# Low level contamination confounds population genomic analysis

**DOI:** 10.1101/2025.01.17.633387

**Authors:** Audrey K. Ward, Eduardo F. C. Scopel, Brent Shuman, Michelle Momany, Douda Bensasson

**Author notes:** These authors contributed equally to this work (Audrey K. Ward, Eduardo F. C. Scopel).

## Abstract

Genome sequence contamination has a variety of causes and can originate from within or between species. Previous research focused primarily on cross-species contamination or on prokaryotes. This paper visualizes B-allele frequency to test for intra-species contamination, and measures its effects on phylogenetic and admixture analysis in two fungal species. Using a standard base calling pipeline, we found that contaminated genomes superficially appeared to produce good quality genome data. Yet as little as 5-10% genome contamination was enough to change phylogenetic tree topologies and make contaminated strains appear as hybrids between lineages (genetically admixed). We recommend the use of B-allele frequency plots to screen genome resequencing data for intra-species contamination.

## Background

The contamination of high-throughput sequence data is a known challenge in genome biology that can lead to incorrect inferences [11, 19, 25, 31]. Low level sample contamination can occur in laboratories during DNA extraction or in culture, at sequencing centers during amplification steps, or even *in silico* if barcodes are not easily distinguished after multiplexing [2, 5, 6, 8]. Most existing tools detect contamination that occurs between species [6]. Yet analysis of bacterial genomes suggests within-species contamination is more likely to lead to mistakes in base calling, species identification or phylogenetic analysis [24]. Furthermore, analysis of RNAseq data for animal mtDNA shows that intra-species contamination can result in the overestimation of heterozygosity and incorrect inference of balancing selection [2].

Most tools for detecting intra-species contamination compare read data to sequence databases for prokaryotes or particular genes or species [6]. A more broadly applicable approach for the detection of within-species contamination is to identify short read data with unusual frequencies of variant alleles after mapping to a reference [8]. A similar approach, the visualization of variant (or B) allele frequencies in plots, is commonly used to determine ploidy or aneuploidy [4, 32]. Using B-allele frequency plots, we encountered low level intra-species contamination in public data for two model fungal species; *Saccharomyces cerevisiae* and *Aspergillus fumigatus*. To determine whether low levels of intra-species contamination are cause for concern, we tested the sensitivity of a standard base calling pipeline, phylogenomic and admixture analyses to within-species contamination using read data that we contaminated *in silico* to known degrees (0 - 50%).

## Results and Discussion

### Within-species contamination in public short read genome data

B-allele frequency plots are routinely used to distinguish homozygous or haploid genome data from heterozygous diploids or polyploids [4, 32]. Single nucleotide polymorphisms (SNPs) in heterozygous diploids differ from the reference genome at read allele frequencies of 1.0 or 0.5, triploids at 1.0, 0.67 or 0.33 and so on (Figure 1). In screening short-read genome data for aneuploidy, we observed public read samples with appreciable levels of intra-species contamination (over 5%) in *S. cerevisiae* (Figure S1, Table S1) and *A. fumigatus* (data not shown). *Saccharomyces cerevisiae* strains are mostly homozygous diploids and *A. fumigatus* strains are usually haploid. We screened *S. cerevisiae* genome data for 1,357 strains sequenced to high read depth (over 30×) and found 8 genomes with at least 5% intra-species contamination [23]. Most of these (N = 6) showed 5 - 10% contamination, and two showed 10 - 20% contamination. Higher levels of contamination would be difficult to distinguish from polyploidy using our methods, but are probably less likely.

**Fig. 1.**
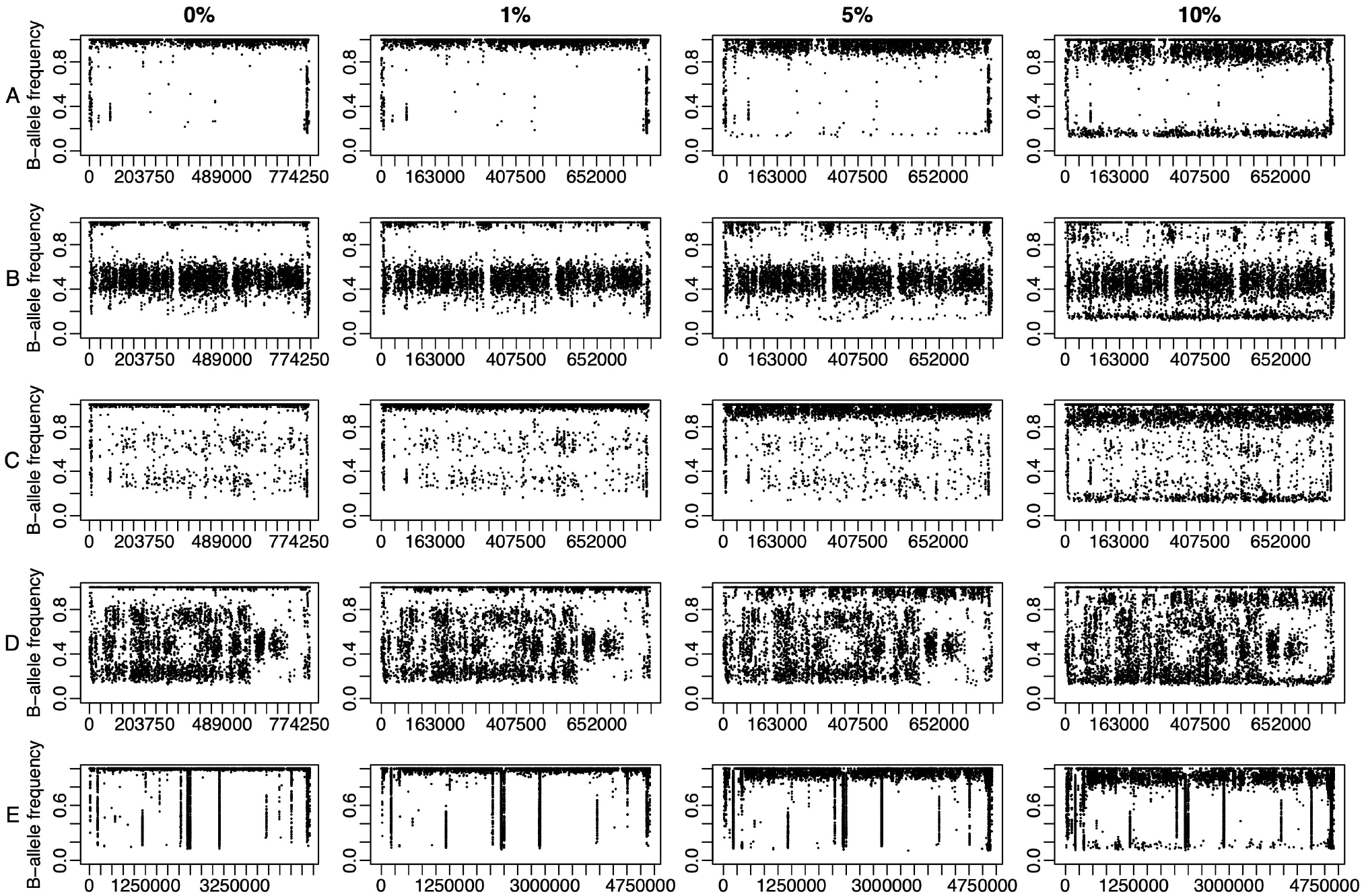
Intra-species contamination is recognizable in B-allele frequency plots at 5% contamination. Plots show base calls for resampled genome data contaminated *in silico* to: 0%, 1%, 5% and 10%. Points show the frequency of non-reference “B” alleles along chromosome II for *S. cerevisiae* (A-D) and along chromosome 1 for *A. fumigatus* (E) for A) a haploid, B) diploid, C) triploid, D) tetraploid and E) haploid. In contaminated mixtures, a substantial fraction of SNP differences from the reference genome appear below their expected frequency of 1.0, at the level expected if the contaminating strain has the same allele as the reference e.g. 0.95 for 5% contamination with a strain matching the reference. In repetitive regions variants appear at many allele frequencies appearing as vertical lines on the plots.

The extent of intra-species contamination in public genome data that we observe for *S. cerevisiae* (0.59%) is lower overall than that reported for bacteria at levels expected to affect base calling (*Escherichia coli* for 0.87%, *Salmonella enterica* 1.48%, *Listeria monocytogenes* 2.22%) [24]. The percentage of *S. cerevisiae* read samples with contamination do however vary greatly by study: from under 0.2% to 15% (Table S1; Fisher’s exact test, *P* = 2 × 10^−6^). This is consistent with past observations that the extent of contamination can differ substantially among studies and sequencing centers [2, 11].

### The effects of *in silico* contamination on base calling

Most contaminated data show only low levels of contamination (5 - 10%; 6 out of 8 contaminated genomes), so correct base calls outnumbered incorrect calls by ten to twenty fold. To determine whether such low level contamination impacts base calling, we examined *in silico* simulations of read data with known levels of added contamination using a standard base calling pipeline. We applied a phred-scaled quality filter (Q40) that labels sites as “low quality” data if they have estimated error rates above 1 in 10,000; a consensus base call would be represented with an “N” and therefore treated as missing data in downstream analyses. The proportion of low quality base calls does not increase with increasing levels of contamination (Table S2). The number of high quality heterozygous base calls does increase with increasing contamination, but in haploids and triploids heterozygosity only reaches the levels seen in diploids and tetraploids with 20% contamination or above. Surprisingly, even the number of high quality homozygous base calls increases slightly at 20% contamination for haploids, and with any amount of contamination at higher ploidy levels (Table S2). These simple quality checks that are easily performed without population genomic analyses suggest that contamination at the levels usually observed in public databases do not greatly affect base calling. However these checks do not address the effects of contamination on variant or SNP sites in particular, which are likely affected differently than invariant sites and are critical for downstream applications.

### Low level contamination affects population genomic analyses

Intra-species contamination likely results in erroneous heterozygous calls at SNP sites. It is therefore not surprising that contamination in past work led to mistakes in estimating the inbreeding statistic, F_*IT*_, which relies on correct heterozygous base calls [2]. Other important population genomic statistics, such as Tajima’s D and ratios of non-synonymous to synonymous diversity were less sensitive to intra-species contamination [2]. The inference of individual ancestry from allele frequency data is useful for estimating population structure and identifying genetic admixture [1]. It also uses heterozygous base calls and is therefore likely sensitive to contamination. Here we tested the effect of contamination at 5% and 10% contamination on the inference of individual ancestry from allele frequency data using the software ADMIXTURE [1]. In all ADMIXTURE runs, 5% contamination did not affect results (Figure S2). The estimation of ancestry was affected however by 10% contamination. In most runs the contaminated strain appeared admixed between donor and recipient lineages and mostly to a greater extent (25%) than the expected 10% contamination level (Figure S3).

In contrast to allele frequency analyses, we expect phylogenetic analyses to be more robust to low levels of contamination because most phylogenetic software treat heterozygous sites as missing data, and we do not expect low level contamination to result in homozygous calls for the minority allele. To test the impact of contamination on phylogenetic analysis, we included a contaminated strain in within-species phylogenomic trees for *A. fumigatus* and *S. cerevisiae*.

Surprisingly, the phylogenetic placement of the recipient strain changed considerably even with only 10% contamination (Figure 2, Figures S4-6). This was true for *S. cerevisiae* and *A. fumigatus* using neighbor joining distance or maximum likelihood approaches. At 10% contamination, we observed major shifts in phylogenetic position (Figures 2, S4-6) and by 20% contamination the recipient *S. cerevisiae* strain clustered with the donor strain (Figures 2 and S4). For *A. fumigatus* we did not include the donor strain in the phylogeny, yet we still saw major changes to tree topology (Figures S5-6). Using neighbor-joining distance, we even saw a small effect on tree topology at 5% contamination in *S. cerevisiae* (Figure 2).

**Fig. 2.**
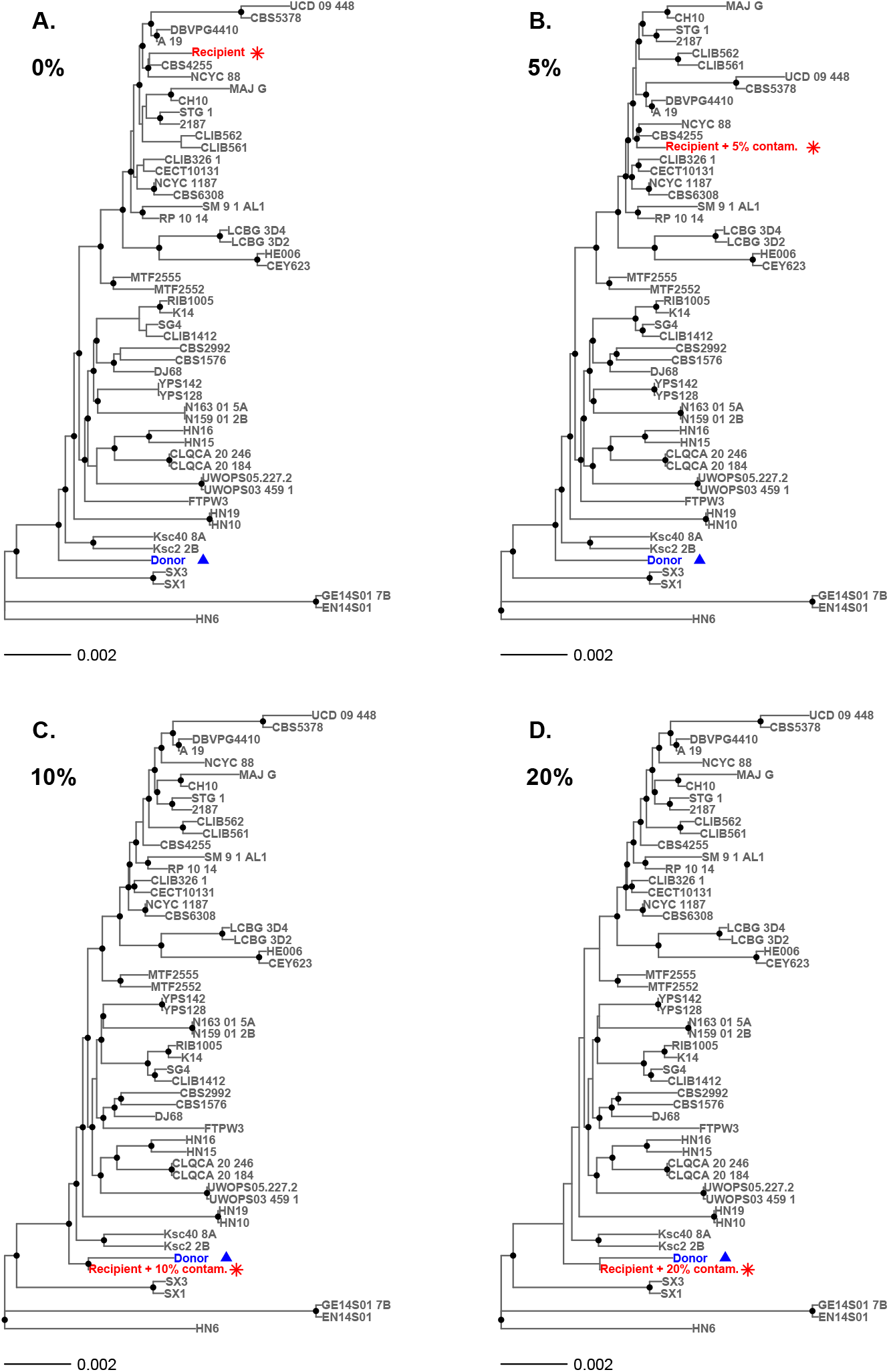
Change in topology for neighbor joining *S. cerevisiae* phylogenetic trees starting at 5% and 10% contamination. Panel A shows donor and recipient genomes in the absence of contamination; B shows the recipient with 5% contamination results in a tree with slightly altered topology; and at higher levels of contamination, 10% in C and 20% in D, the recipient clusters with the donor strain.

How could low level (5-10%) contamination alter tree topology? In contaminated data, differences between the donor and recipient sequence appear as heterozygous sites (Table S2) [2]. These sites no longer contribute to estimates of genetic distance between the contaminated strain and donor lineage using most phylogenetic software [17]. In addition, the donor alleles will be called in regions where the recipient genome has low quality sequence or deletions relative to the reference, which could explain the increase in homozygous base calls at increasing levels of contamination (Table S2). In cases where the donor genome has more high quality regions mapping to the reference than the recipient genome (as in this study; Table S2) enough homozygous base calls might result from donor reads to change tree topology. The chances of seeing an effect of contamination on cluster analyses also increase with increasing divergence between donor and recipient genomes [24]. The strains we used in this study are from genetically distinct lineages (Figures 2, S2-6), so our analyses probably represent a worst-case scenario.

## Conclusions

Here we show that within-species contamination of genome data can lead to incorrect phylogenies or inference of genetic admixture, even at the low levels seen in public databases. Contamination has led to incorrect conclusions in the past, but most reports are on between-species contamination or Sanger sequencing studies [11, 19, 25, 31]. Analysis of intra-species contamination in bacteria show that it can be especially damaging [24]. Using eukaryotic models, we show the importance of screening for intra-species contamination in short read genome data, especially because phylogenetic analyses can be more sensitive to contamination (5-10%, Figure 2) than previously recognized (40-50%) [24]. The visualization of SNPs in mapped read data with B-allele frequency plots provides a means to synchronously assess ploidy, heterozygosity and potential contamination (Figure 1); all important information for downstream phylogenetic or population genomic analyses.

## Methods

To understand the effects of intra-species contamination on base calls and phylogenomic analysis, we created contaminated mixtures with various levels of contamination for *A. fumigatus* and *S. cerevisiae*; 0, 1, 5, 10, 20, 30, 40, and 50%. Using published short-read data [12, 28] (Table S3), *S. cerevisiae* haploid, heterozygous diploid, triploid, and tetraploid genomes (CBS1479, DBVPG1074, NPA05a1, UCD 06-645) were contaminated with reads from a haploid donor (CLIB219.2b). For *A. fumigatus*, the recipient and donor were haploid strains eAF749 and eAF163 respectively. The reads for each mixture were randomly sampled without replacement using seqtk sample (v1.2 for *S. cerevisiae*; 1.3 for *A. fumigatus*; https://github.com/lh3/seqtk). For each *S. cerevisiae* strain (12 Mbp genome), we used 8 million paired reads; 4 million for each simulated fastq file. For each *A. fumigatus* (29 Mbp genome) we used 6 million reads; 3 million for each simulated fastq file.

For base calling, we mapped reads to reference genomes using Burrows-Wheeler Aligner (bwa mem, v0.7.17; [15]). The reference genomes were SacCer_Apr2011/sacCer3 from strain S288c at UCSC for *S. cerevisiae* and ASM265v1 from strain Af293 for *A. fumigatus* [20]. Consensus sequences were generated using SAMtools mpileup and BCFtools call -c (v1.6) [16] with indels removed and read depth limited to a maximum of 100,000 reads. Mapped alignments were converted to fasta format using vcfutils.pl vcf2fq (from BCFtools) and seqtk seq with a phred-scaled quality threshold of 40 to define low quality base calls. Mitochondrial DNA was removed for downstream analyses. We generated BAF plots using vcf2alleleplot.pl with default options and counted high quality heterozygous and homozygous sites in fasta files using basecomp.pl [3].

To test the effects of contamination on phylogenetic analyses, we compared each recipient or contaminated genome to a panel of reference strains with known phylogenetic positions (Table S3). For *S. cerevisiae*, we randomly selected up to 2 strains (where available) from each of the known 26 lineages described in Scopel *et al*. [28], which resulted in a total of 52 reference panel strains including the donor strain. Recent genetic admixture is common in *S. cerevisiae* [18] and can complicate phylogenetic analysis, but prior analyses show that none of the strains used here were admixed [22, 28]. For *A. fumigatus*, we used one-dimensional k-means clustering to categorize 168 strains [12] into 52 clusters based on their pairwise genetic distances from a single strain (CF098), then randomly chose a single strain from each cluster. Genetic distances were estimated using the dnadist function of PHYLIP (v3.697) with default parameters and a 0.5:1 transition:transversion ratio [9] and we used python to perform the k-means clustering (getGenDist.py; [27]). Neighbor-joining trees were constructed using MEGA (v10.0.5) [14] with the Tamura-Nei model [30] and 100 bootstrap replicates. Maximum likelihood trees were estimated using RAxML (v8.2.11 for *S. cerevisiae* and 8.2.12 for *A. fumigatus*) [29] with a GTRGAMMA model and 100 bootstrap replicates. For visualization, trees were rooted with EN14S01, GE14S01 7B and HN6 for *S. cerevisiae* and JN10 for *A. fumigatus* then right ladderized using the ape package (v5.8; [21]) in R (v4.3.3).

For analysis of population structure and genetic admixture in *S. cerevisiae*, we used ADMIXTURE (v1.3.0) [1]. Each recipient or contaminated genome was merged into an alignment with the sequence of 52 reference panel strains using BCFtools view with the –min-ac 1 option (v1.15.1). Low-quality reads (phred score under 40) were filtered in VCFtools (v0.1.16) [7]. Alignment files were converted to text and binary files using PLINK (v1.9b 6.21) [26]. Genomes were assigned to populations (genetic clusters) in repeated ADMIXTURE runs with default parameters and varying numbers of populations (K); from 2 to 26 with five replicates per K. Resultant ancestry proportions were aligned across K values using CLUMPAK distruct (v1.1) [13] and results were visualized using the R package pophelper (v2.3.1) [10]. The Fisher’s exact test and other analyses and visualizations were performed in R (v4.3.3).

## Supplementary information

Supplementary information accompanies this paper. **Additional file 1: Table S1**. Differences among studies in rates of intra-species *Saccharomyces cerevisiae* contamination. **Table S2**. Intra-species contamination does not lower the quality of base calls. **Figure S1**. B-allele frequency plots show intraspecies contamination in public *S. cerevisiae* genome data. **Figure S2**. No effect of 5% contamination on analysis of genetic admixture. **Figure S3**. Contamination levels of 10% result in incorrect calls of genetic admixture. **Figure S4**. Change in topology for neighbor joining *A. fumigatus* phylogenetic trees with 10% contamination. **Figure S5**. Change in topology for maximum likelihood *S. cerevisiae* phylogenetic trees with 10% contamination. **Figure S6**. Change in topology for maximum likelihood *A. fumigatus* phylogenetic trees with 10% contamination. **Additional file 2: Table S3**. Summary of the strains used for *in silico* contamination and population genomic analyses.

## Acknowledgements

We would like to thank Jacqueline Peña for help with data management ahead of publication, and Momany and Bensasson lab members for helpful discussion.

## Declarations

### Ethics approval and consent to participate

Not applicable

### Consent for publication

Not applicable

### Availability of data and materials

SRA accession numbers for the data used are listed in Additional file 2: Table S3. The code developed in this project, getGenDist.py, is available on GitHub and archived at Zenodo with doi 10.5281/zenodo.13881957.

### Authors’ contributions

All authors conceived and designed the experiment. EFCS screened for contamination in public data, developed the *in silico* sampling approach and performed base call analyses; AKW performed the phylogenetic and admixture analyses; BS performed all analyses for *A. fumigatus*. All authors wrote the manuscript.

