## Supplemental Figures and Tables S1-S2 for "Low level contamination confounds population genomic analysis"

### Supplemental Tables

**Table S1** Differences among studies in rates of intra-species *Saccharomyces cerevisiae* contamination.

| SRA identifier | Uncontaminated | Contaminated | % | Study |
|---|---|---|---|---|
| ERP014555 | 915 | 0 | 0% | [1] |
| PRJNA396809 | 260 | 5 | 2% | [2] |
| PRJEB7601 | 55 | 0 | 0% | [3] |
| PRJNA1090965 | 43 | 0 | 0% | [4] |
| PRJEB11698 | 17 | 3 | 15% | [5] |

Data from Peña et al [4] after excluding genomes with low read depth and studies with fewer than 10 high depth genomes. Rates appear different (Fisher's exact test, $P = 2 \times 10^{-6}$) even after excluding PRJEB11698 (Fisher's exact test, $P = 0.002$).

**Table S2** Intra-species contamination does not lower the quality of base calls

| % from donor.[1] | Ploidy | Homozygous | Heterozygous | Low Quality | P(LQ)[2] |
|---|---|---|---|---|---|
| *A. fumigatus* | | | | | |
| 100% | haploid | 28,056,212 | 27,124 | 1,299,087 | 0.044 |
| 0% | haploid | 27,191,111 | 26,332 | 2,166,779 | 0.074 |
| 1% | | 27,194,419 | 26,678 | 2,163,182 | 0.074 |
| 5% | | 27,303,362 | 30,663 | 2,050,218 | 0.070 |
| 10% | | 27,616,509 | 44,955 | 1,722,747 | 0.059 |
| 20% | | 28,018,909 | 75,478 | 1,289,836 | 0.044 |
| 30% | | 28,104,794 | 83,560 | 1,195,866 | 0.041 |
| *S. cerevisiae* | | | | | |
| 100% | haploid | 11,461,857 | 3,697 | 604,481 | 0.050 |
| 0% | haploid | 11,430,299 | 2,706 | 638,192 | 0.053 |
| 1% | | 11,432,806 | 2,784 | 635,631 | 0.053 |
| 5% | | 11,448,751 | 4,868 | 617,607 | 0.051 |
| 10% | | 11,456,000 | 24,042 | 591,174 | 0.049 |
| 20% | | 11,464,068 | 72,671 | 534,541 | 0.044 |
| 30% | | 11,471,388 | 80,901 | 519,008 | 0.043 |
| 40% | | 11,473,538 | 81,923 | 515,830 | 0.043 |
| 50% | | 11,474,291 | 82,281 | 514,720 | 0.043 |
| 0% | diploid | 11,426,767 | 62,889 | 581,602 | 0.048 |
| 1% | | 11,427,772 | 62,895 | 580,591 | 0.048 |
| 5% | | 11,431,346 | 64,035 | 575,878 | 0.048 |
| 10% | | 11,430,347 | 76,131 | 564,783 | 0.047 |
| 20% | | 11,435,185 | 104,413 | 531,692 | 0.044 |
| 30% | | 11,439,882 | 108,814 | 522,575 | 0.043 |
| 40% | | 11,441,299 | 108,547 | 521,427 | 0.043 |
| 50% | | 11,441,050 | 105,473 | 524,656 | 0.043 |
| 0% | triploid | 11,421,825 | 8,610 | 637,335 | 0.053 |
| 1% | | 11,423,877 | 8,665 | 635,663 | 0.053 |
| 5% | | 11,425,285 | 10,582 | 631,913 | 0.052 |
| 10% | | 11,415,094 | 28,950 | 624,236 | 0.052 |
| 20% | | 11,413,683 | 75,713 | 579,163 | 0.048 |
| 30% | | 11,419,975 | 83,775 | 564,165 | 0.047 |
| 40% | | 11,423,533 | 84,218 | 559,800 | 0.046 |
| 50% | | 11,423,917 | 83,808 | 560,476 | 0.046 |
| 0% | tetraploid | 11,347,790 | 46,526 | 676,765 | 0.056 |
| 1% | | 11,349,837 | 46,590 | 674,801 | 0.056 |
| 5% | | 11,376,595 | 47,656 | 646,986 | 0.054 |
| 10% | | 11,414,485 | 59,758 | 596,994 | 0.049 |
| 20% | | 11,439,265 | 96,163 | 535,856 | 0.044 |
| 30% | | 11,446,409 | 101,228 | 523,665 | 0.043 |
| 40% | | 11,449,966 | 98,988 | 522,346 | 0.043 |
| 50% | | 11,452,171 | 95,042 | 524,088 | 0.043 |

[1]Percent of reads from the donor (contaminant) genome. The recipient genome has 0% of reads from donor. For *Aspergillus fumigatus*, the donor strain was eAF163 and the recipient was eAF749. For *Saccharomyces cerevisiae*, the donor strain was CLIB219.2b and haploid, heterozygous diploid, triploid, and tetraploid recipient strains were CBS1479, DBVPG1074, NPA05a1, UCD_06-645 respectively. Donor and recipient genomes were resampled to the same read depth as contaminated genomes.

[2]Proportion of base calls that are low quality (phred-scaled quality lower than Q40).

#### Supplemental Figures

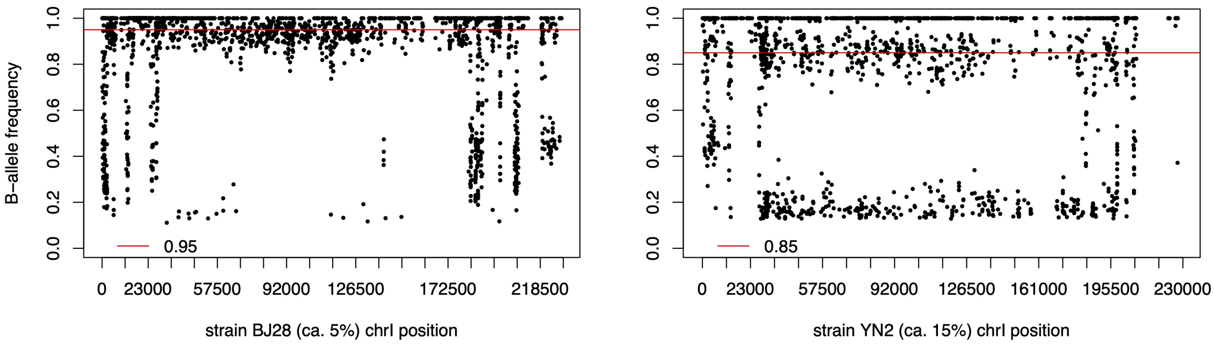

**Figure S1** B-allele frequency plots show intraspecies contamination in public *S. cerevisiae* genome data. In the left panel, the points clustering around the red line at 0.95 suggest contamination levels of 5% where the sequenced strain (BJ28) has SNPs that differ from the reference strain, but the contaminating strain allele matches the reference. The right panel shows B-allele frequencies for strain YN2 suggesting 15% contamination.

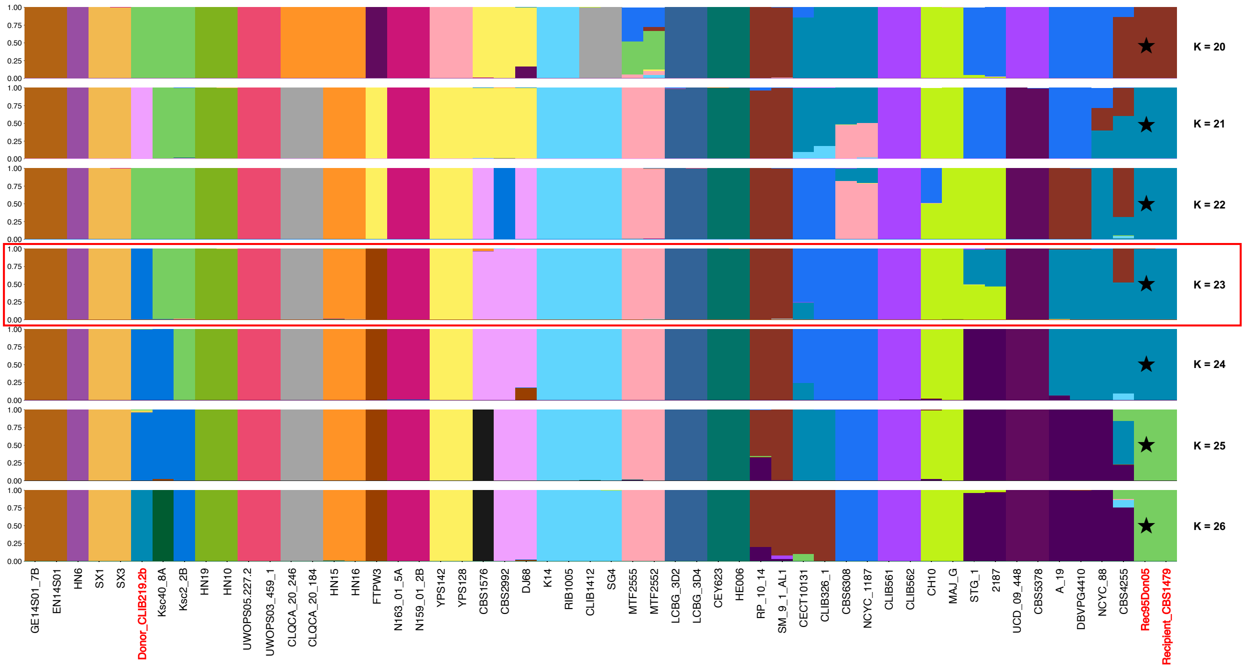

**Figure S2** No effect of 5% contamination on analysis of genetic admixture. The plot shows the ancestry proportions for each individual arranged in the order seen in Figure S5. These plots show the runs with the highest log-likelihoods for each assumed number of populations (K = 20 - 26), and the run with the most clustering similarity to the phylogenetic analyses (K = 23) is highlighted with a red box. Individual genomes highlighted with red text are the donor genome (Donor_CLIB10_2B), and the recipient genome showing the ancestry proportions expected with 0% contamination (Recipient_CBS1479) and an *in silico* mix of recipient with 5% contamination from the donor (Rec95Don05, black star) showing the same results as the uncontaminated recipient.

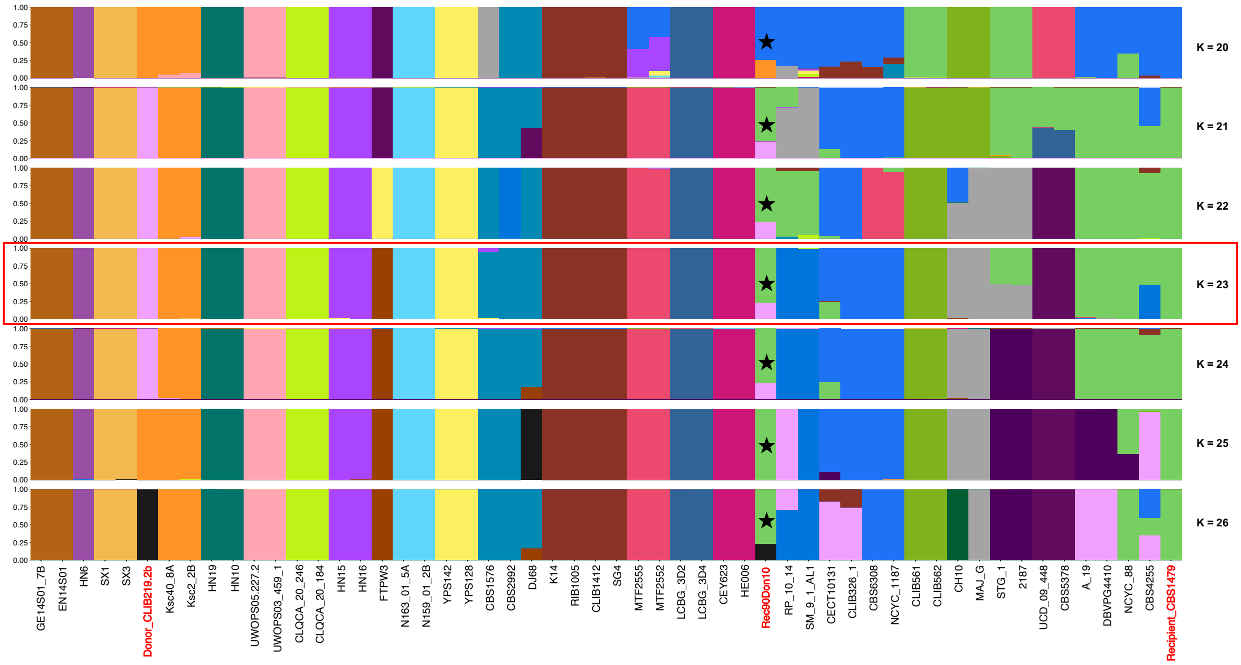

**Figure S3** Contamination levels of 10% result in incorrect calls of genetic admixture. The plot shows the ancestry proportions for each individual arranged in the order seen in maximum likelihood phylogenetic analyses Figure S5. These plots show the runs with the highest log-likelihoods for each assumed number of populations (K = 20 - 26), and the run with the most clustering similarity to the phylogenetic analyses (K = 23) is highlighted with a red box. Individual genomes highlighted with red text are the donor genome (Donor_CLIB10_2B), and the recipient genome showing the ancestry proportions expected with 0% contamination (Recipient_CBS1479) and an *in silico* mix of recipient with 10% contamination from the donor (Rec90Don10, black star). In most runs, the genome with 10% contamination shows 25% admixture between donor and recipient lineages.

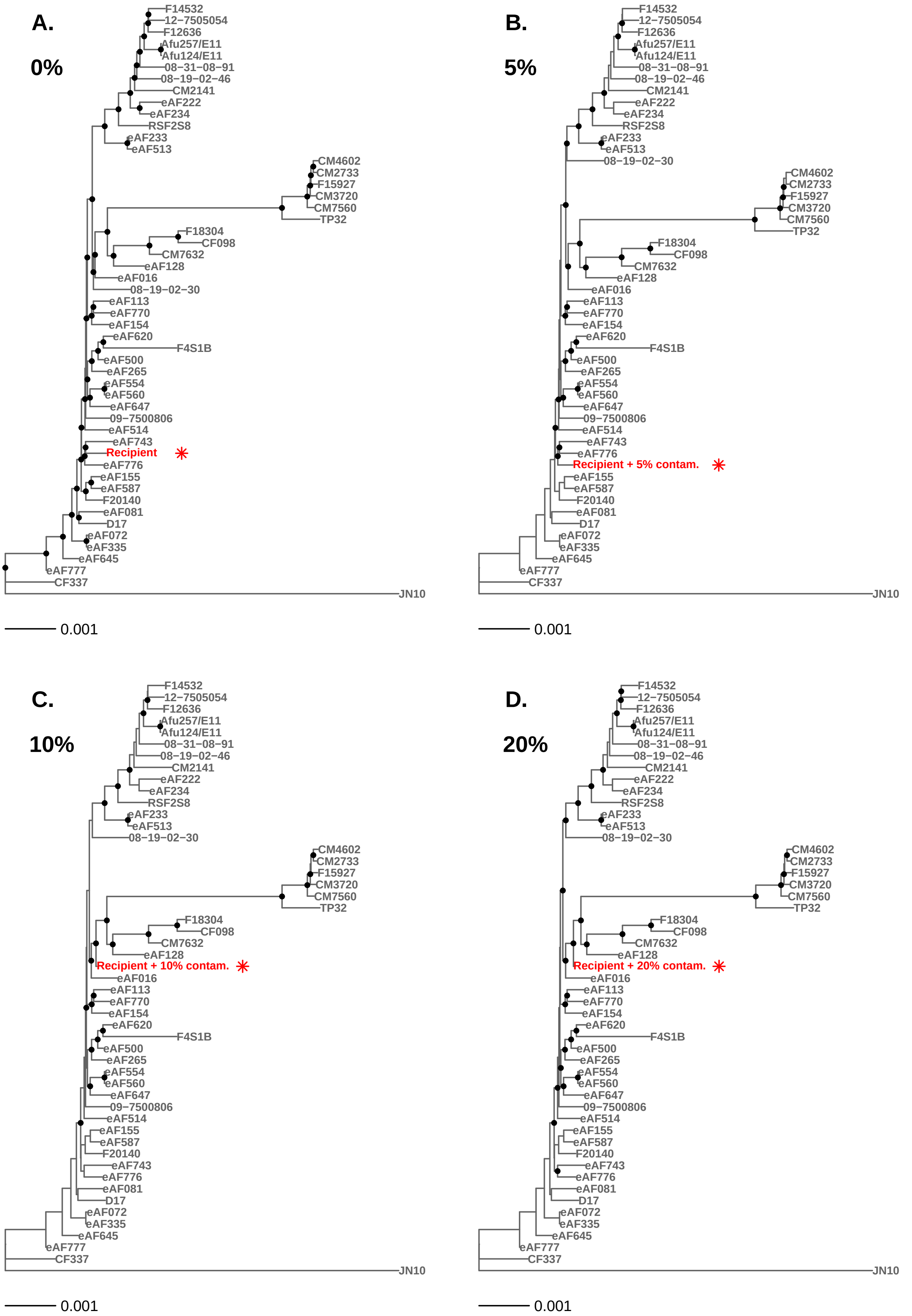

**Figure S4** Change in topology for neighbor joining *A. fumigatus* phylogenetic trees with 10% cross-contamination. Panel A shows the recipient genome in the absence of cross-contamination; B shows the recipient with 5% contamination; C shows a new topology at 10% contamination and D shows the same new topology at 20%.

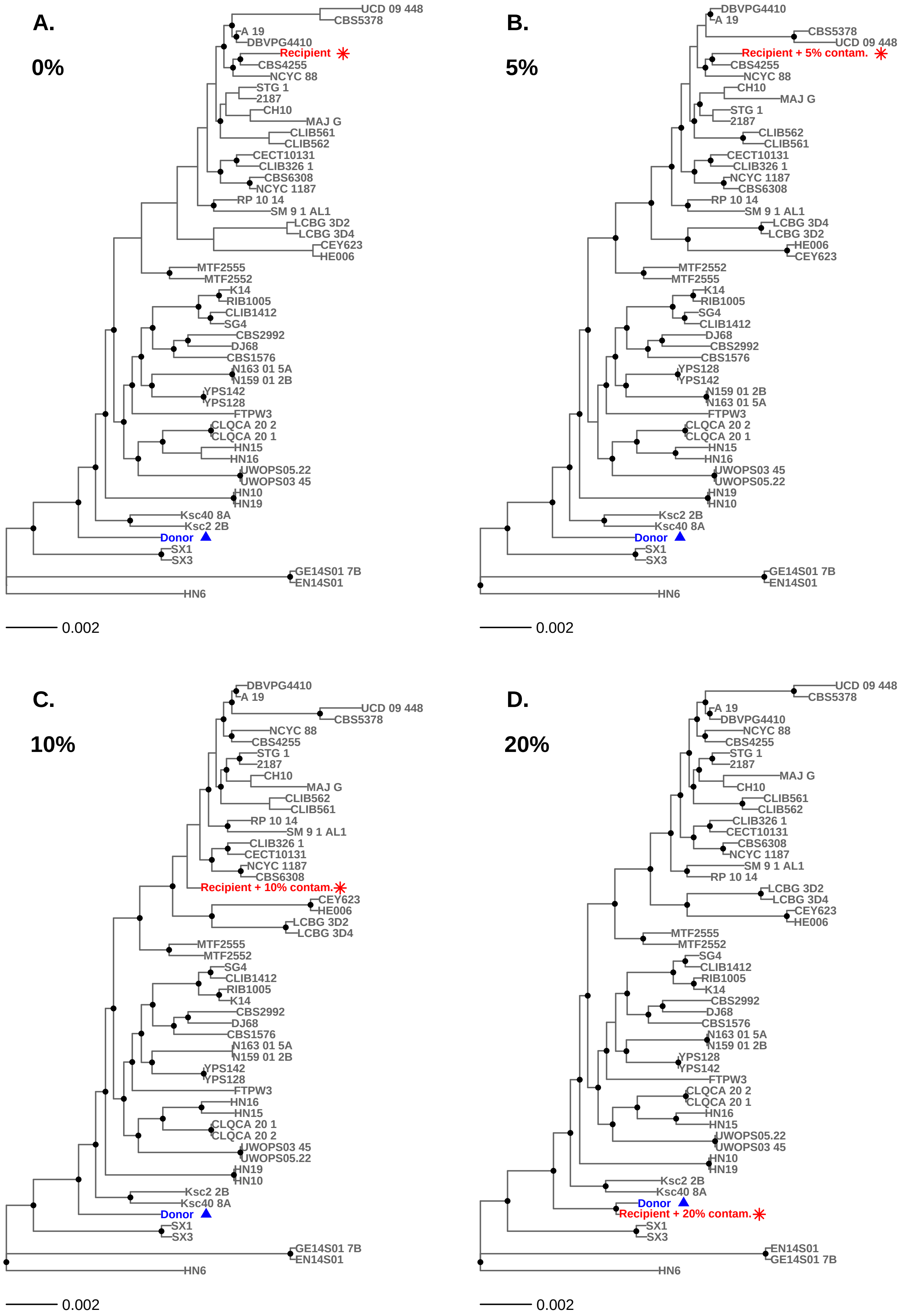

**Figure S5** Change in topology for maximum likelihood *S. cerevisiae* phylogenetic trees with 10% cross-contamination. Panel A shows donor and recipient genomes in the absence of cross-contamination; B shows the recipient with 5%; C shows a new topology at 10%; D shows a more greatly altered topology at 20% contamination.

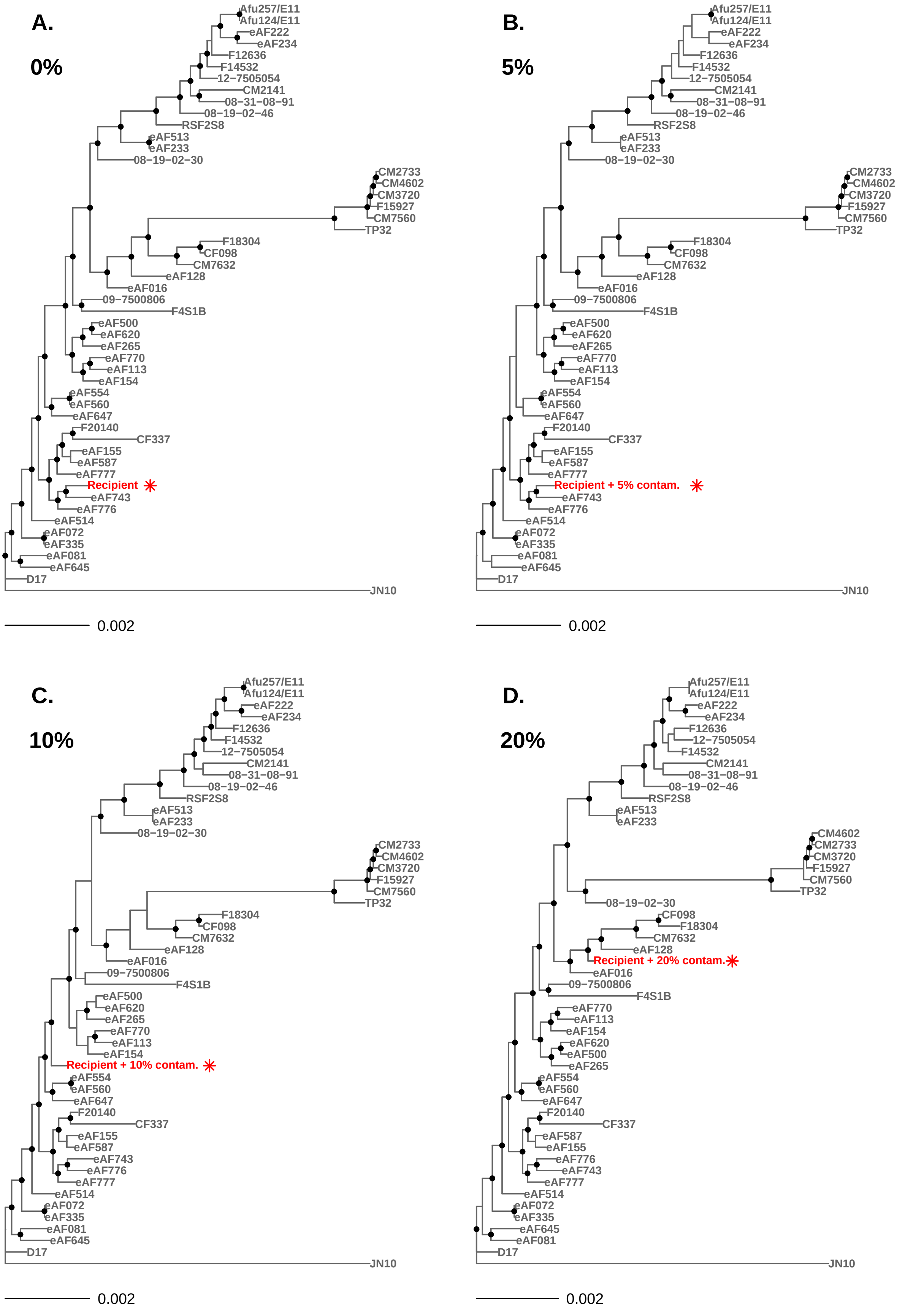

**Figure S6** Change in topology for maximum likelihood *A. fumigatus* phylogenetic trees with 10% cross-contamination. Panel A shows the recipient genome in the absence of cross-contamination; B shows the recipient with 5%; C shows a new topology at 10%; D shows a more greatly altered topology at 20% contamination.
